# The crowns have eyes: Multiple opsins found in the eyes of the Crown-of-Thorns Starfish *Acanthaster planci* including the first r-opsin utilized by a deuterostome eye

**DOI:** 10.1101/173187

**Authors:** Elijah K. Lowe, Anders Garm, Esther Ullrich-Lüter, M. Ina Arnone

## Abstract

Opsins are G protein-coupled receptors used for both visual and non-visual photoreception, and these proteins evolutionarily date back to the base of the bilaterians. In the current sequencing age, phylogenomic analysis has proven to be a powerful tool, facilitating the increase in knowledge about diversity within the opsin subclasses and, so far, nine paralogs have been identified. Within echinoderms, opsins have been studied in Echinoidea and Ophiuroidea, which do not possess proper image forming eyes, but rather widely dispersed dermal photoreceptors. However, most species of Asteroidea, the starfish, possess true eyes and studying them will shed light on the diversity of opsin usage within echinoderms and help resolve the evolutionary history of opsins. Using high-throughput RNA sequencing, we have sequenced and analyzed the transcriptomes of different *Acanthaster planci* tissue samples: eyes, radial nerve, tube feet and a mixture of tissues from other organs. At least ten opsins were identified, and eight of them were found significantly differentially expressed in both eyes and radial nerve, providing new important insight into the involvement of opsins in visual and nonvisual photoreception. Of relevance, we found the first evidence of an r-opsin photopigment expressed in a well developed visual eye in a deuterostome animal.

## Introduction

Light carries an immense amount of information about the surroundings. Direct light from the sun, the moon, or the stars is used by various animals to set diurnal or annual clocks and to set direction during navigational tasks. Light reflected from the surroundings guides innumerous different behaviours as it provides information about objects with unprecedented details and speed. Light reception is thus widespread in the animal kingdom but interestingly the molecular machinery behind light reception shares many common features across all phyla. In most cases examined so far the first step in the phototransduction, the absorption of the photons in metazoan, is mediated by a specific protein family called opsins [1]. Opsins are seven transmembrane G protein-coupled receptors binding a chromophore, retinal, which undergoes a conformational change upon the absorption of light, thus triggering the rest of the transduction cascade. During the last couple of decades, several molecular studies have examined the diversity of the opsin family and found nine major clades [2]. What the work has also shown is that many animals have a surprisingly high number of opsin gene copies and that they can be expressed in almost any body region or organ [3]. In many of these cases, the functions remain unknown and may well be outside light reception [4,5].

Light reception is known from all major groups of echinoderms and is facilitated by different types of photoreceptors ranging from non-pigmented dermal photoreceptors to proper image forming eyes. A rather special case has been suggested in the brittlestar *Ophiocoma* with dispersed microlenses formed by skeletal elements [6]. Another dermal photoreceptor system is found in sea urchins which has also been suggested to support image forming vision [7,8]. The genome has been sequenced for the sea urchin *Strongylocentrotus purpuratus* and eight opsin genes were found belonging to the opsin clades c-opsins, r-opsins, Go-opsins, peropsin, neuropsin and echinopsins A and B [9,10]. The latter two groups were recently renamed as bathyopsins and chaopsins, respectively, [2]. The r-opsin Sp-opsin4 is expressed in cells at the base of the transparent tube feet and is putatively mediating the directional negative phototaxis described for a couple of species [8,11,12]. The brittle star *Amphiura filiformis* has even higher opsin diversity with at least 13 gene copies [13], but here little is known about expression patterns and behavioural roles.

Dermal photoreception is also known from several species of starfish (Asteroidea) [14-16] but it is unknown if it is opsin based and if so whether it is the same opsin as found in the photoreceptors of the eyes. Remarkably, these echinoderms also have well defined eyes. Those eyes are found in most non-burrowing starfish species at the tip of each arm sitting at the base of the unpaired terminal tube foot as a direct extension of the radial nerve. They are compound eyes and structurally they resemble the eyes of arch clams and fan worms [17,18] with lensless ommatidia typically 20-40 μm in diameter. Depending on species, adult specimens have 50-300 ommatidia in each eye and recent studies have shown that this supports spatial resolution in the range of 8-17 degrees used for navigation [19-21]. These studies have also indicated that the ommatidia have a single population of photoreceptors which utilize an opsin with peak sensitivity in the deep blue part of the spectrum around 470 nm.

Recently, two draft genomes of the crown-of-thorns starfish (COTS) *Acanthaster planci*^1^, relative to animals collected from Okinawa, Japan and Great Barrier Reef, Australia (GBR), were released. These two genomes shared 98.8% nucleotide identity and were determined to be the same species [22], which is in agreement with previous classification of the Pacific ocean COTS being one species [23]. Although *A. planci* is not the first Asteroidea with an assembled genome, it is the first species with well defined eyes to have an assembled genome. The GBR genome was released along with annotations for ~24,500 protein coding genes [22], and this presents an opportunity to study the mechanisms behind vision in a species with a well defined eye that is evolutionarily close to species with alternative methods of photoreception. Here we have used tissue specific transcriptomics to investigate the differential expression of opsin genes in *A. planci*. We found at least seven different paralogs and, by comparing expression levels in the eyes, in locomotory tube feet, in the radial nerve, and in a mixture of gonadal, stomach, and epidermal tissue, we are able to infer which opsins are most likely used in vision, in non-visual photoreception, and outside photoreception.

## Material and methods

### (a) Animals

The specimens of *A. planci* used in this study were hand collected on the Great Barrier Reef off the coast of Cairns, Australia. After collection the animals were kept in holding tanks with running seawater at 26 degrees for 2-3 days and then flown to Denmark. In Denmark they were kept under similar conditions at the Danish National Aquarium, The Blue Planet, where they were fed three times a week with a past of enriched squid and fish meat. Tissue samples were taken from four specimens with diameters of 15-23 cm. Three terminal tube feet including the eye, 3 locomotory tube feet, approx. 5 cm radial nerve, and pieces of the gonads, the stomas, and the epidermis were sampled from each of the four specimens and stored in RNAlater at 4ºC. Two additional animals were collected at the coastal reefs of Guam and a total of 12 eyes and 10 cm radial nerve were taken from them directly after collection and stored in RNAlater at 4ºC.

### (b) RNA extraction and sequencing

The tissue samples were removed from the RNAlater, immediately frozen with liquid nitrogen and homogenized using a mortar and pestle. Powdered tissues were then dissolved in EuroGOLD RNAPure (EMR 506100) and processed using EUROzol RNA extraction protocol (EMR055100, euroclone), then subjected to LitCl (4M) purification. Library preps and sequencing were done at Università degli Studi di Salerno using SMART-Seq v4 Ultra Low Input RNA Kit.

Sequenced reads were examined using fastqc and then quality filtered and trimmed using trimmomatic (v0.33) [25]. Quality controlled reads were quasi-mapped and quantified to v1 great barrier reef *Acanthaster planci* transcriptome using salmon (v0.82) [26]. Transcripts per million (TPM), the normalized transcript counts [27]. Differentially expressed genes were identified using DESeq2 [28] (FDR ≤ 0.05 and -1.5 ≥ log2FC ≥ 1.5). All scripts can be found at https://github.com/elijahlowe/Acanthaster_opsins.git.

### (c) Identification of opsin sequences

Opsin protein sequences were collected from echinoderms [35] (Freeman et al. 2008), molluscs [39,40], vertebrates [49], and NCBI [50], as well as from publications themselves, as described in electronic supplementary material (SI 1). These sequences were used to perform Reciprocal Best Hits (RBH) BLAST against the *A. planci* (GBR) proteome. Additionally, pantherSCORE2.0 [51] was used to identify opsin sequences using hidden Markov models. The collected sequences were aligned with MAFFT (v7.215) [52,53] using L-INS-i algorithm which is better designed for divergence sequences and performed well when benchmarked against other multiple sequence aligners [54]. The aligned sequences were then trimmed using trimAl [55], removing gaps that occurred in 10% of the alignments while being sure to retain 60% of the total sequence length. Initial phylogenetic trees were generated using the aligned sequences with fasttree (v2.1.7) [56] and visualised with figtree (v1.4.3) [57]. Additional trees were then generated using iqtree [58] with 10,000 ultrafast bootstrap support [59] using the ‘a Bayesian-like transformation of aLRT’ (abayes) method [60], and LG+F+R6 amino acid substitution model selected using ModelFinder [61]. Modifications such as additional labels and visual effects were done using inkscape.

## Results

### Diversity of opsins in *A. planci*

We identified thirteen putative opsin paralogs in *A. planci*’s proteome using Reciprocal Best Hits (RBH) BLAST and pantherSCORE2.0, which share high sequence similarity. Closer examination using the genome revealed that two of these sequences were actually one fragmented opsin (gbr.65.47.t1 and gbr.65.48.t1), so we manually edited the sequence, which can be found in the opsin fasta file of electronic supplementary material (SI 2). These sequences were included in a phylogenetic analysis totaling 169 sequences spanning 40 different species. Our phylogenetic analysis from both FastTree and iqtree revealed ten *A. planci* opsins belonging to 7 groups and two sequences clustering with the melatonin receptors outgroup (figure 1): 1 rhabdomeric opsin (r-opsin), 4 ciliary opsins (c-opsins), 1 peropsin, 1 Go-opsin, 1 RGR opsin, 1 neuropsin, and 1 chaopsin. Chaopsins were first classified as echinoderm specific [2]. *A. planci* possesses representatives of all so far known echinoderm opsin groups with the exception of bathyopsin (former echinopsin A), which thus has yet to be identified in any starfish. *A. planci* opsins grouped closest with those of *Patiria miniata-an* eyeless starfish-followed by *Asteria rubens* opsins.

**Figure 1.**
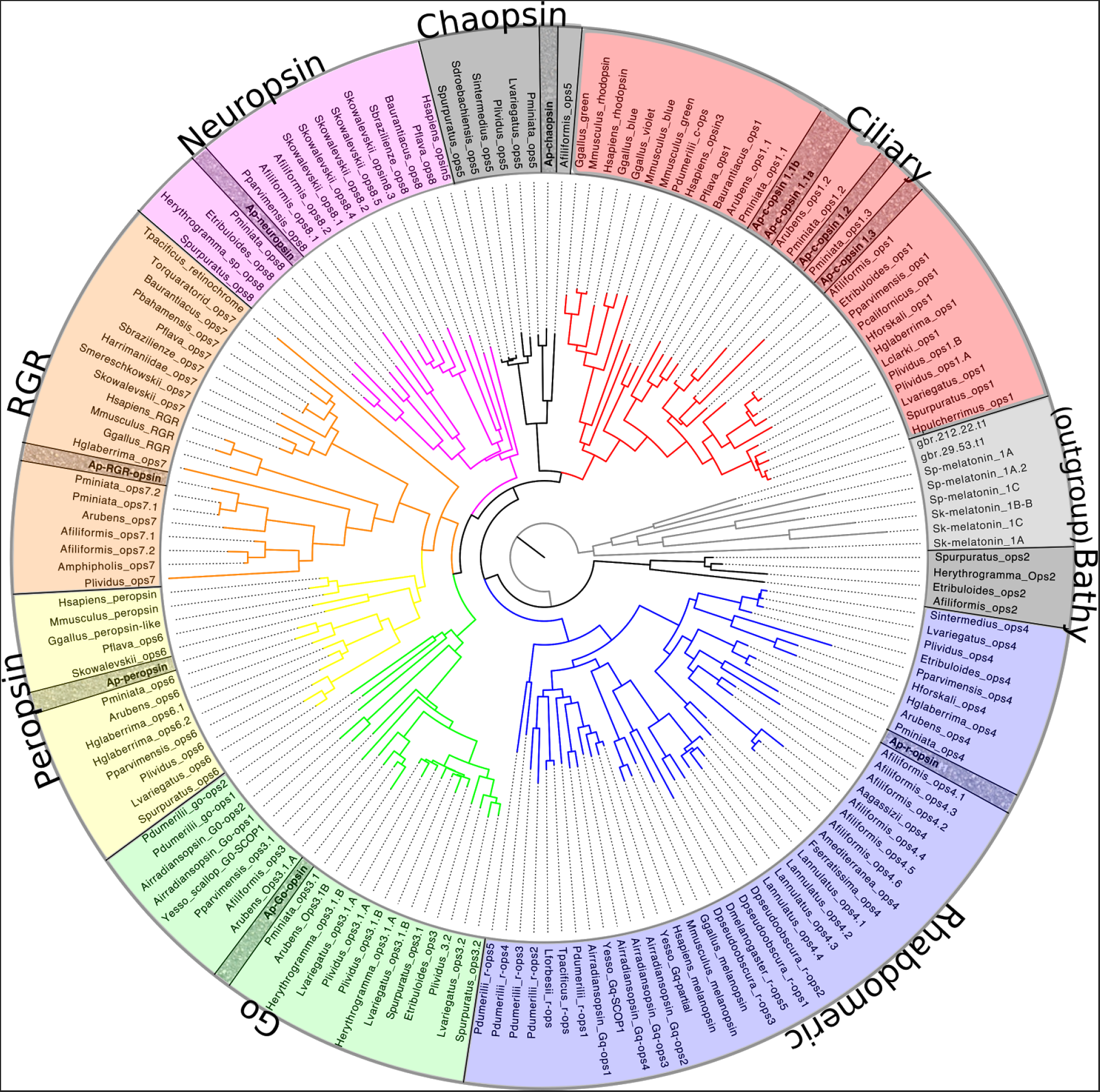
Phylogenomic tree of 169 opsin sequences with melatonin receptors as the outgroup. There are 10 *A. planci* opsins (bold and checkered background) which classify into 7 different groups: 4 c-opsins, 1 chaopsin, 1 r-opsin, 1 peropsin, 1 RGR opsin, 1 go-opsin, and 1 neuropsin. No bathyopsin was found in *A. planci*, and has yet to be identified in any starfish species. The tree was generated by iqtree [58] with 10,000 ultrafast bootstrap support [59] using the ‘a Bayesian-like transformation of aLRT’ (abayes) method [60], and LG+F+R6 amino acid substitution model.

In *A. planci* eyes, all opsins with the exception of ciliary opsin 1.1b and neuropsin were up-regulated in comparison to the mixed tissues (Figure 2). This was also the case when comparing opsin expression in the radial nerve to the mixed sample but to a lesser degree (Figure S1). Expression of opsins in the tube feet, however, was not observed to be up-regulated compared to the mixed tissue, in fact c-opsin 1.1b and neuropsin showed higher expression in the mixed tissue samples (figure 3).

**Figure 2.**
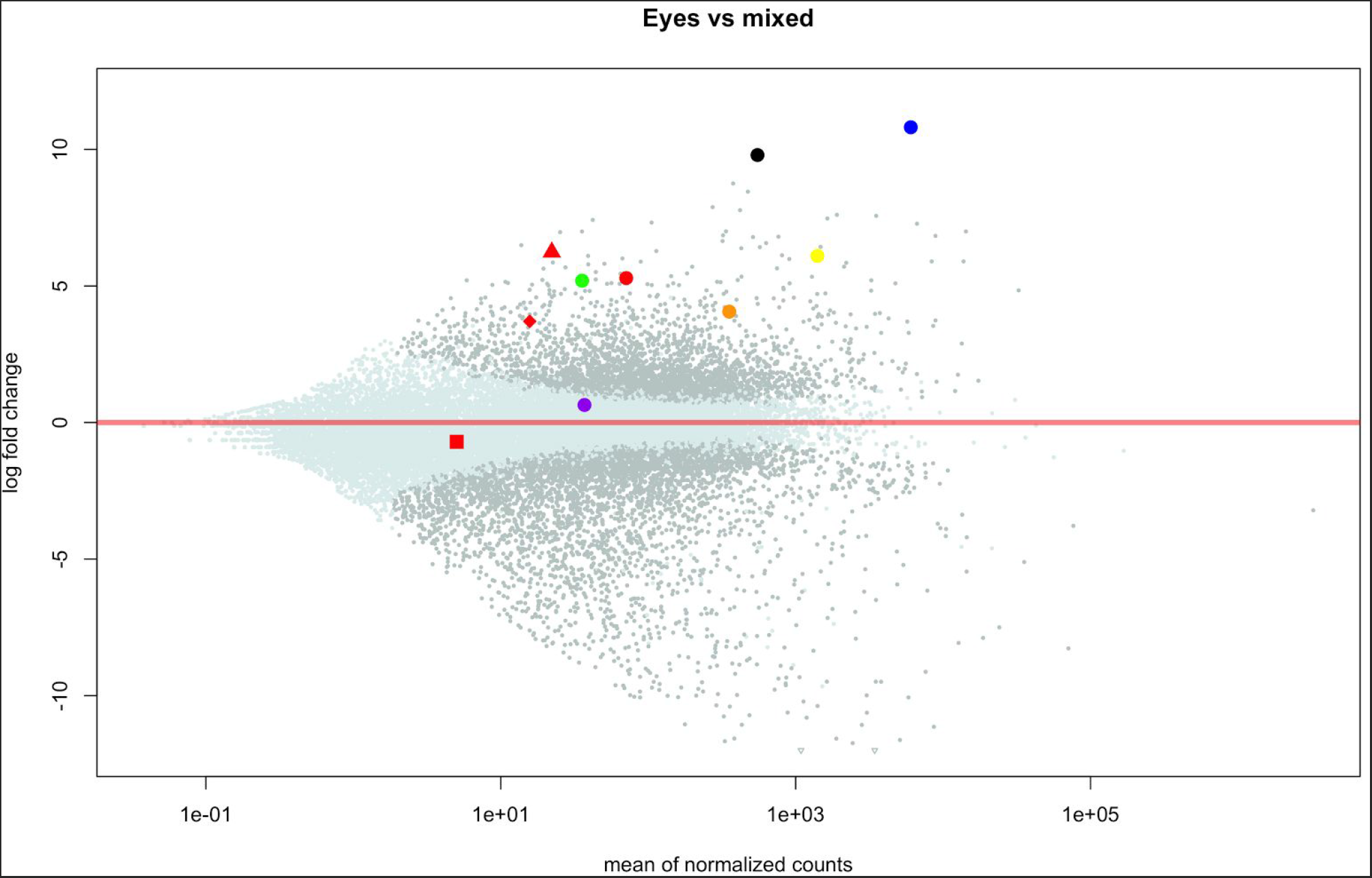
Summary of differential expression data in various *A. planci* tissue samples focusing on opsin expression. (A) Transcripts per million (TPM) counts of each opsin for each tissue type. For eyes (green) the highest expressed opsins are r-opsin, peropsin, chaopsin, RGR opsin, and go-opsin, with c-opsin 1.2, c-opsin 1.3, and neuropsin being expressed at low amounts, while no expression is observed in c-opsin 1.1b. RGR opsin and peropsin were the highest expressed amongst the other tissues. The mixed tissue and tube feet (tf) have little to no expression (TPM ≤ 0.5) of c-opsin 1.2, c-opsin 1.3, go-opsin, and chaopsin.

**Figure 3.**
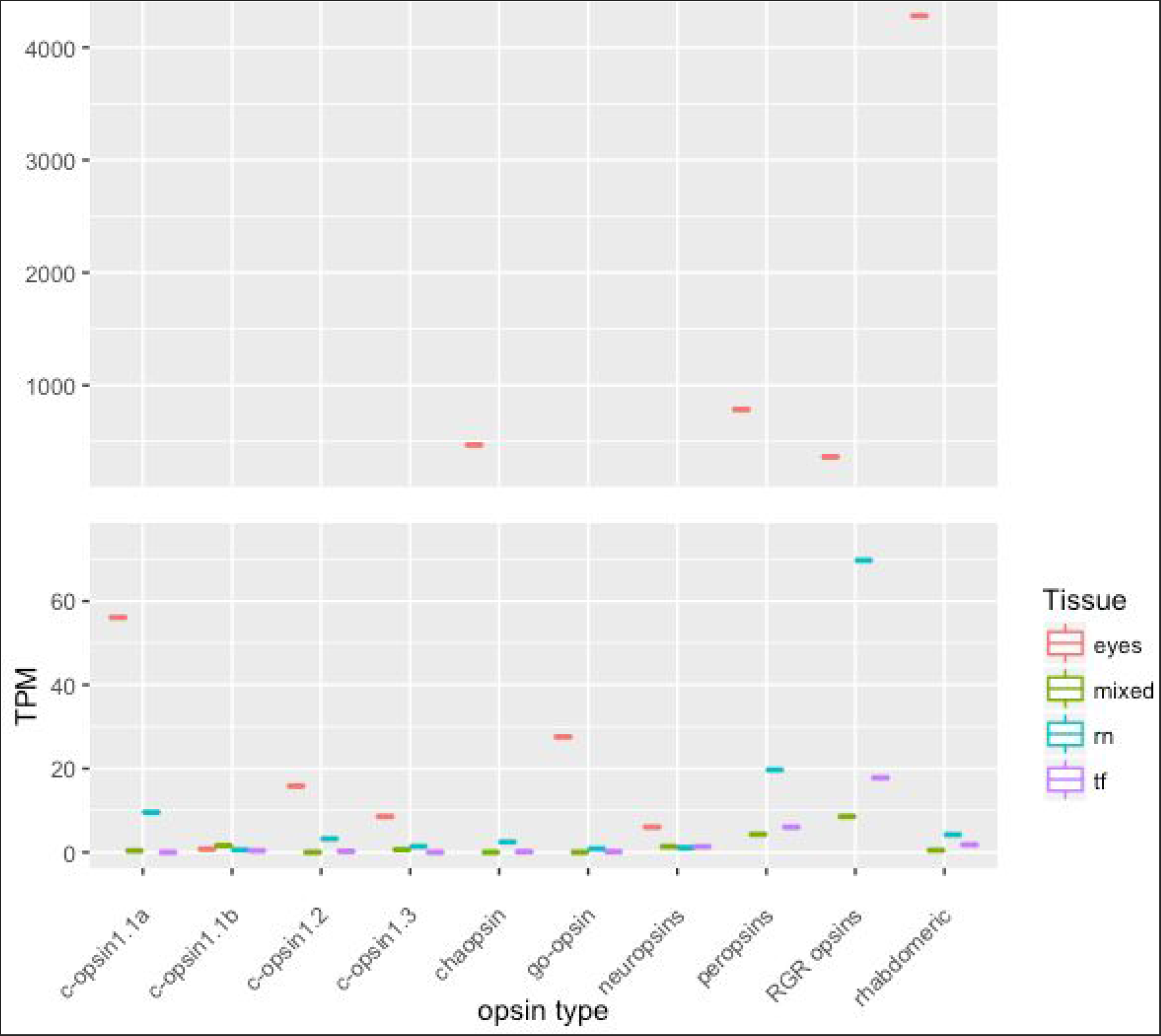
Differential expression of the eye tissue samples versus the mixed tissue samples. R-opsin is the most differentially expressed, followed by chaopsin. With the exception of c-opsin 1.1b and neuropsin, all opsins are differentially expressed in the eyes of *A. planci* compared to the mixed tissue samples.

In order to assess the putative functionality of the *A. planci* identified opsin sequences, an analysis of the key residues necessary for opsin function was performed (Table 1). In most opsins, the retinal binds to the K296 via a Schiff base bond, however the proton in the opsin protein is unstable and a counterion is needed and often supplied by the highly conserved Glutamic acid (E113) residue. There are cases, however, where this residue is replaced by a Tyrosine (Y), Phenylalanine (F), Methionine (M), or Histidine (H) and the other highly conserved Glutamic acid residue, E181, serves as the counterion [62]. This is the case with the majority of the opsins in *A. planci*, where E113 are replaced with a Tyrosine (Y), with the exception of Ap-Go-opsin which has a Isoleucine (I) in the position 113.

**Table 1.** Analysis of known typically highly conserved key residues and amino acid motifs

| *A. planci* Opsin | Disulfide bond<sup>a</sup><br>(C110/C187) | LxxxD<sup>b</sup><br>(79) | Counterion<sup>c</sup><br>(E113/E181) | Schiff base<sup>d</sup><br>(K296) | NPxxY<sup>e</sup><br>(302) |
| --- | --- | --- | --- | --- | --- |
| C-opsin 1.1a<br>(gbr.65.47.t1) | C/C | ICVAD | Y/E | K | NPVIY |
| C-opsin 1.1b<br>(gbr.508.2.t1) | C/C | VCVAD | Y/- | K | NPVIY |
| C-opsin 1.2<br>(gbr.65.46.t1) | C/F | ISVGD | Y/- | R | N---- |
| C-opsin 1.3<br>(gbr.65.45.t1) | L/C | VCEA- | Y/A | K | NPIIY |
| Go-opsin<br>(gbr.470.6.t1) | C/C | MAVSD | I/E | K | NPLIY |
| R-opsin<br>(gbr.176.5.t1) | C/C | LAFSD | Y/E | K | NPLVY |
| Chaopsin<br>(gbr.176.10.t1) | C/C | LSGSD | Y/E | K | NPIIY |
| Peropsin<br>(gbr.31.87.t1) | C/C | ASAGD | Y/E | K | NPLMF |
| RGR opsin<br>(gbr.37.113.t1) | C/C | LCAGD | Y/E | K | NAALQ |
| Neuropsin<br>(gbr.37.57.t1) | C/C | LAVSD | Y/E | K | NPIIY |

a. Motif required for recognition of rhodopsin by G-protein [81]

b. Motif interacting with NPxxY motif upon receptor activation for structural constra[82]

c. Glutamic acid residues stabilizing the Schiff base bond

d. Lysine residue forming Shiff base bond with retinal Motif providing structural constraints in response to photoisomerization during formation of the G protein-activating Meta II [83]

In addition to ten opsin sequences, we have also observed ten *A. planci* sequences that are potential G protein alpha subunits. Phylogenomic methods classified these sequences as 3 Gα_s_, 1 Gα_o_, 4 Gα_i_, 1 Gα_q_, and 1 Gα_12_ (figure S2). All identified G protein alpha subunits with the exception of 1 Gα_s_ (gbr.231.19.t1), 1 Gasα_i_ (gbr.143.10.t1) and the Gα_12_ are upregulated in the eyes of *A. planci* compared to the mixed tissue samples (figure S3).

#### (a) Ciliary and rhabdomeric opsins

There are four ciliary opsins identified in the *A. planci* genome, three of which are expressed in the eyes; c-opsin 1.1a, 1.2 and c-opsin 1.3. These three c-opsins were closely clustered on gbr_scaffold65 (~70 kb), and observed to be significantly differentially expressed in the animal, as neither are expressed in the mixed tissue (figure 4). Of the Ap-c-opsins, 1.1a is the highest expressed, followed by Ap-c-opsin 1.2 and 1.3. C-opsin 1.2 was observed to not have the K296 residue, which is required for the formation of the Schiff base, but instead an Arginine (R) residue is placed in this position. This is the only *A. planci* opsin missing this key residue. While the E113 counterion is replaced with tyrosine (Y) in all of the Ap-c-opsins, the E181 counterion is present in Ap-c-opsin1.1a, completely missing in Ap-c-opsin 1.1b and 1.2 and replaced with an Alanine (A) in Ap-c-opsin 1.3 The Ap-c-opsin 1.2 and Ap-c-opsin 1.3 are missing other important motifs, the C187 and C110 disulfide bond motifs [63], respectively (table 1).

**Figure 4.**
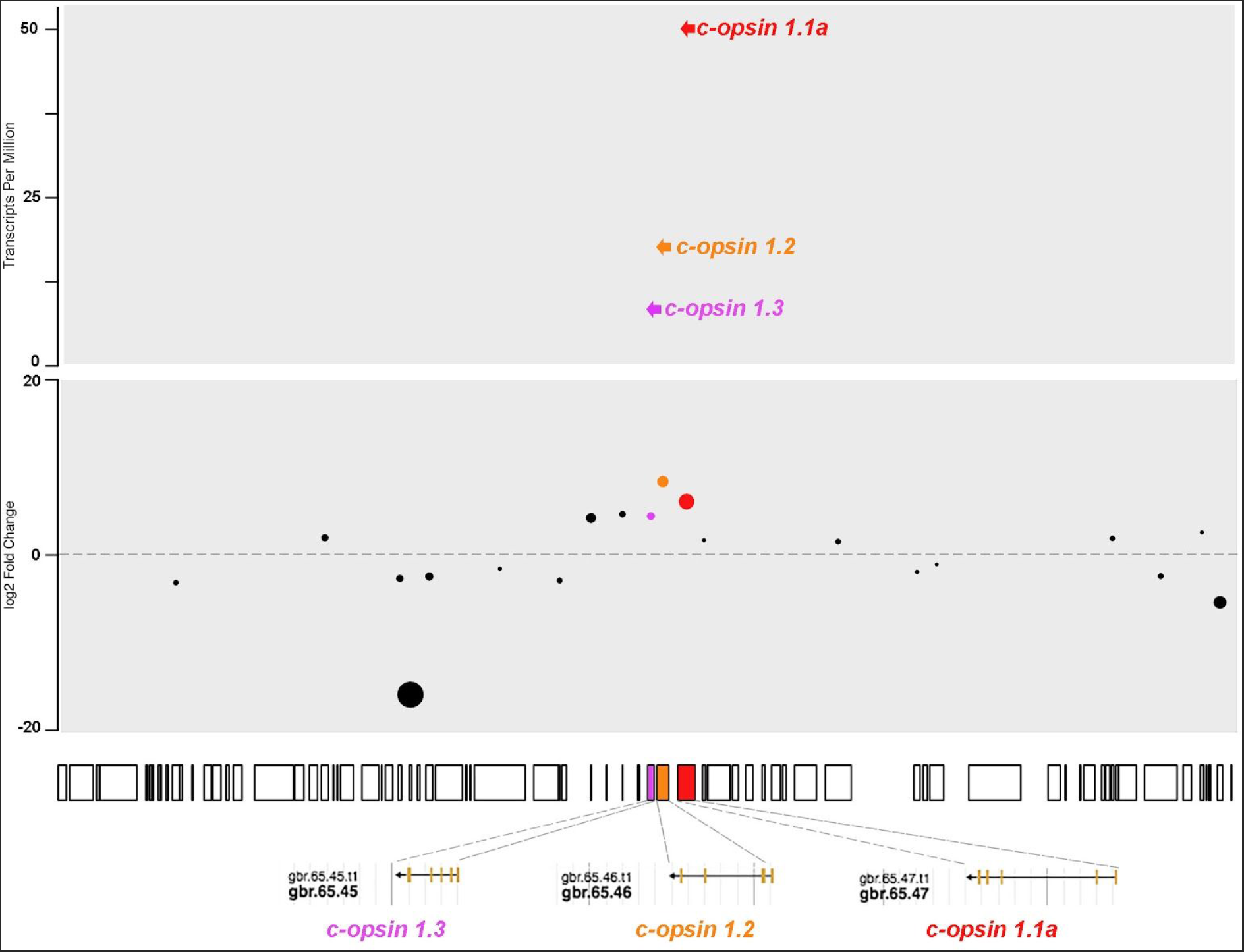
The boxes below the two panels represent “gbr_scaffold65” from the Great barrier reef assembly of the *A. planci* genome. Each box represents a gene and the size of the box correlates the to size of the gene. Three of the four Ap-c-opsins are located on the same scaffold all being transcripted 3’ to 5’; Ap-c-opsin 1.1a (red), Ap-c-opsin 1.2 (orange) and Ap-c-opsin 1.3 (purple). Bottom panel: The log2 fold change of significantly differentially expressed genes (FDR ≤ 0.05) comparing *A. planci* eyes the mixed tissue samples, with the three Ap-c-opsins highlighted. Top panel: Expression of the three Ap-c-opsins clustered on “gbr_scaffold65”, the expression level correlates with the spatial ordering within the cluster (in the direction 3' to 5'), the most 3’ having the highest expression and the most 5’ having the lowest expression.

*A. planci*’s r-opsin, on the other hand, is the most highly differentially expressed of the opsins and of any other genes when comparing eye tissues to mixed tissue (figure 2). Further, the Ap-r-opsin was observed to be the most up-regulated in the starfish eye tissue and its sequence features the Lysine residue (K296), critical for the Schiff base formation, and a putative counterion (E181) (Table 1).

#### (b) Chaopsin

Chaopsin is a recently identified group of opsins. Ramirez et al. [2] found the formerly described groups of anthozoa I opsins [64] and the echinoderm echinopsin B [10] to cluster forming the chaopsin group. In agreement with D’Aniello et al. (2015) we found an *A. planci*’s chaopsin to cluster together with other ambulacrarian chaopsins between the r-opsin and c-opsin clades (figure 1). Ap-chaopsin is amongst the highest differentially expressed opsins in our *A. planci* transcriptomes, with ~9.7 log_2_ fold changes in the eye compared to the mixed tissue (figure 2). It is also expressed in the radial nerve tissue, but to a far lesser degree, and appears to not be significantly expressed in the mixed tissues or the tube feet (figure 3 and S1).

#### (c) Peropsin RGR and Go opsins

Peropsin and RGR opsin are the highest expressed of the opsin in the tube feet, mixed, and radial nerve, although both still have higher expression in the eyes (figure 3 and S1). The disulfide bond linkage C110/C187, counterion sites and the Lysine for the Schiff base formation are all present. However, both *A. planci* RGR-opsin and peropsin contain variations of the NPxxY motif, NAALQ and NPLMF, respectively (Table 1). Additionaly, peropsin has a variation of the LxxxD motif, ASAGD. Ap-Go-opsin was expressed in both the eyes and radial nerve but was not found in the mixed sample or the tube feet (figure 2 and 3).

## Discussion

Light sensing is an important aspect of life and much of it is mediated or initiated by the G-protein-coupled receptor proteins, opsins. The release of the annotated draft genome of *A. planci* has prompted us to investigate its opsin repertoire and expression in a tissue specific manner. This allowed us to classify the specific opsins and to infer possible function and further expand the knowledge of opsin evolution especially within deuterostomes. Ten opsins were identified spanning seven clades: r-opsin, c-opsin, Go-opsin, peropsin, neuropsin, RGR opsin and chaopsin. Opsins have also been sequenced from two other starfish species, *Asterias rubens* and *Patiria miniata*. Through a phylogenomic analysis, it was observed that *A. planci* opsins grouped closest to those in the eyeless *P. miniata*. This grouping is in accordance with the phylogenetic position of these starfish species [65], with *A. planci* as an Acanthasteridae more closely related to *P. miniata*, an Asterinidae, (both species belonging to Valvatida)- than to *A. rubens* belonging to Forcipulatacea. However, studies on tissue specific opsin expression is needed to reveal in which organs of the eyeless starfish *P. miniata* the respective opsin orthologs are expressed. To this point, it remains unclear if opsins potentially serving a visual function in the eye possessing *A. planci* might have switched functions in the eyeless representative, or if their expression simply persists as a potential evolutionary remain, a finding known e.g. from blind cave salamanders which still express opsins inside their highly degenerated and pigmentless photoreceptors [66].

### Rhabdomeric opsin

Electrophysiological recordings from the photoreceptors of *A. planci* have shown light absorption here to be utilizing a single opsin. Of the several opsins found to be expressed in the eyes of *A. planci*, r-opsin appeared as the highest differentially expressed gene in the eye transcriptome when compared to mixed samples, thus suggesting this to be the opsin utilized for vision. A similar function has been proposed for the sea urchin r-opsin in *S. purpuratus* tube feet [67]), no deuterostome eye has previously been found to express an r-opsin. As *A. planci* eyes have been previously demonstrated to perform proper spatial vision [19-21], our findings thus provide first evidence for a deuterostome eye utilizing an r-opsin for spatial vision.

### Ciliary opsin

Ciliary opsins are well known for their role in vertebrate vision and brain function in some invertebrates [68]. [69]). Clustering of opsins has so far not been described outside deuterostomes, but it is also seen in the echinoderm, *S. purpuratus* where Go-opsins (opsin 3.1 and 3.2) clustered on scaffold26 (Echinobase; [49]). The significance of the occurrence of opsin clustering in echinoderms is not obvious and awaits more functional studies.

### Chaopsin

Chaopsin (Ap-chaopsin) is the second most differentially expressed opsin in *A. planci* eyes and it has many of the motifs necessary for phototransduction, including the NPxxY binding motif in the 7 transmembrane domain involved in coupling with the G protein. Little is known about the functions of chaopsins (or opsin5) but they have been identified in Echinoidea, Asteroidea, and Ophiuroidea [30] and *Strongylocentrotus intermedius* [70]. In the Caribbean elkhorn coral, *Acropora palmata*, opsin3, which along with echinoderm opsin5 belongs to the chaopsin clade [71]. This leads us to hypothesize that the Ap-chaopsin may be important for phototransduction in *A. planci* and potentially in all echinoderms. The exact role of this opsin remains elusive, though.

### Peropsin and RGR opsin

Peropsin and RGR opsin were expressed in the tube feet, the mixed tissue, and the radial nerve of *A. planci*, but both have the highest expression in the eyes. They are considered as photoisomerase enzymes and not as photopigments, since they bind to all-trans retinaldehyde to regenerate 11-cis-retinoids for pigment regeneration. This has been observed for peropsin in the vertebrate retinal pigment epithelium [72], in cephalopod photoreceptors [73] and in the jumping spider [74]. Knock-down mice [75], together with biochemical and spectroscopic studies in amphioxus [76], have demonstrated the same properties for RGR opsin in chordates. This family of opsins is thus important for visual pigment regeneration [77]. While RGR opsins are known to not have the NPxxY binding motif in the 7 transmembrane domain involved in coupling with the G protein, this motif is often found in peropsins [46].

### Go-opsin

In *A. planci* eye and radial nerve transcriptomes, we found significant expression of a Go-opsin along with putative Go alpha subunit proteins. Evidence for Go-opsins in animals is rare. These opsins interact with a specific G-protein that differs from those in the c-opsin as well as the r-opsin transduction cascade [79]. Knocking down Go-opsin did not lead to absence of phototaxis in *P. dumerilii* but did reduce the sensitivity to the blue-cyan part of the color spectrum (λ_max_=488 nm). A similar absorbance spectrum (λ_max_=483 nm) was observed in amphioxus, *Branchinostoma belcheri*, Go-opsin [76,80]. While the morphological and expression data on Go-opsin in *P. yessonensis* point towards a functioning in a visual context, no other studies have found this. Our data does not reveal if *A. planci* Go-opsin is co-expressed with any other opsins inside the same cells, nonetheless, the fact that this opsin is expressed in the starfish eye along with the results from annelids and amphioxus, opens up the possibility that it is involved in spectral tuning of vision in *A. planci*. Such a tuning would indeed be consisted with the spectral sensitivity curves obtained by electrophysiology in this starfish species [20].

## Conclusion

In conclusion, our findings demonstrate that the eye of the starfish *A. planci* expresses an entire set of ten different opsin proteins. This starfish has recently been demonstrated to perform spatial vision in order to prey on its preferred coral food and its r-opsin, as the by far most highly expressed photopigment in its eyes, is likely to facilitate these sophisticated visually guided behaviours. *A. planci* is thus not only one more echinoderm possessing a much more complex photoreceptor system than previously assumed, but rather the first deuterostome animal that has been shown to navigate by the use of r-opsin expressing photoreceptors. The variety of opsins found differentially expressed in various starfish light sensitive tissues by our transcriptomic analyses also set the groundwork for comparative studies on evolutionary changes in photoreceptor function that occurred towards the vertebrate eye.

## Data accessibility

Raw sequencing data will be submitted to NCBI SRA under the accession number SUB2926139. Additionally, fasta files containing all sequences can be found in the data section of the github repository https://github.com/elijahlowe/Acanthaster_opsins and in electronic supplementary material.

## Authors' contributions

Collection of tissue samples were done by AG. RNA extraction was performed by (MIA) and EKL. Computational analysis was performed by EKL. Writing of the paper was performed by EKL, AG, IA and EUL. All authors contributed to the project design.

## Acknowledgement

We would like to thank Ronald Petie for help collecting the animals and Jérôme Delroisse for his expertise and assistance in reviewing the manuscripts. Additionally, we would like to take Michigan State University for the use of their High Performance Computing Cluster (HPCC).

---

**Figure S1.**
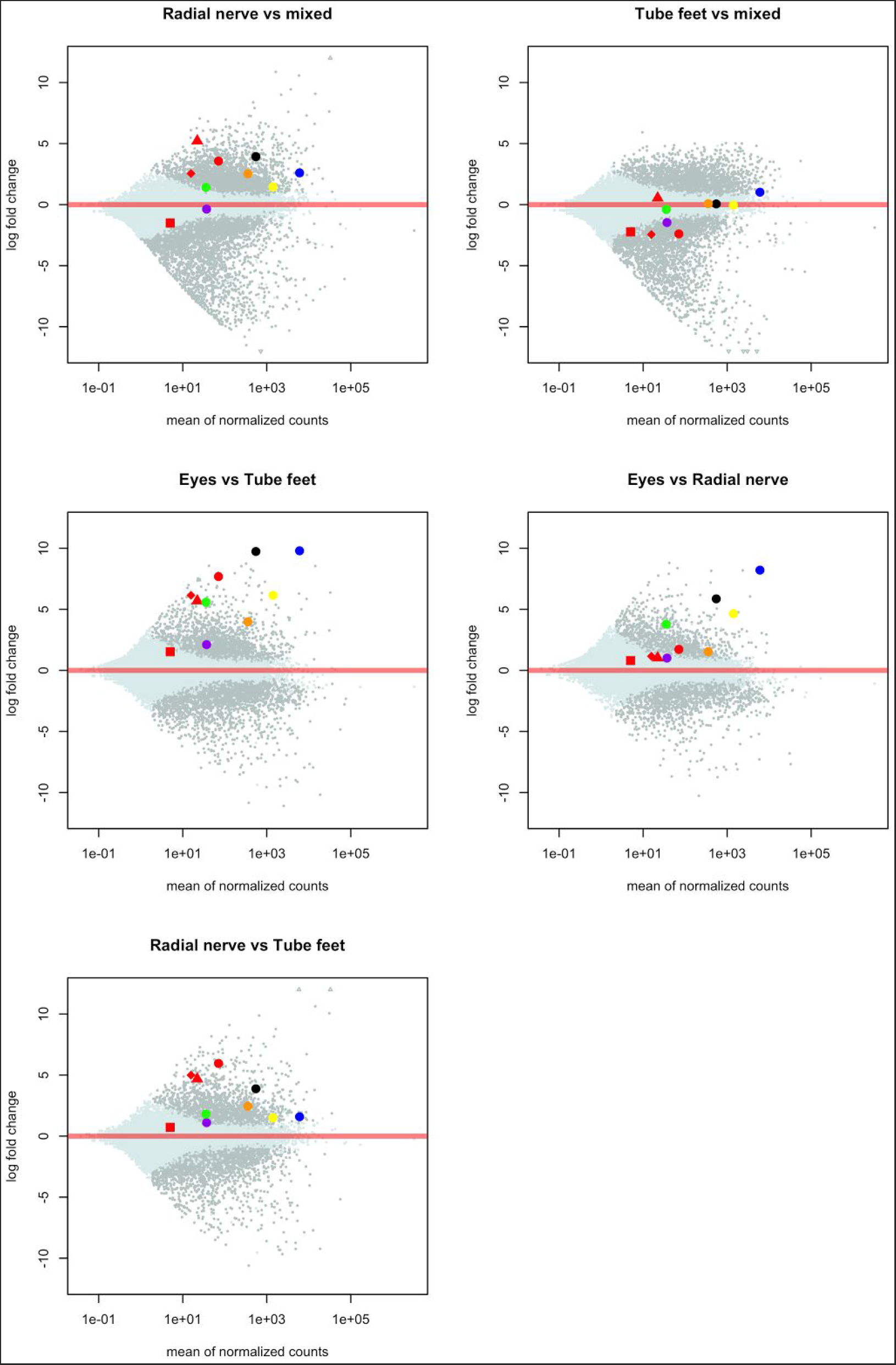
Differential gene expression in all *A. planci* tissue samples with the opsins highlighted: c-opsins in red, go-opsins in green, chaopin in black, neuropsin in purple, peropsin in yellow, r-opsin in blue and RGR opsin in orange. The y-axis in the log_2_fold-change, as the distance from the y-axis increases the more differentially expressed a gene is in one tissue versus the other. The x-axis represents counts per million (CPM), an increase on the this axis shows genes with more reads counts.

**Figure S2.**
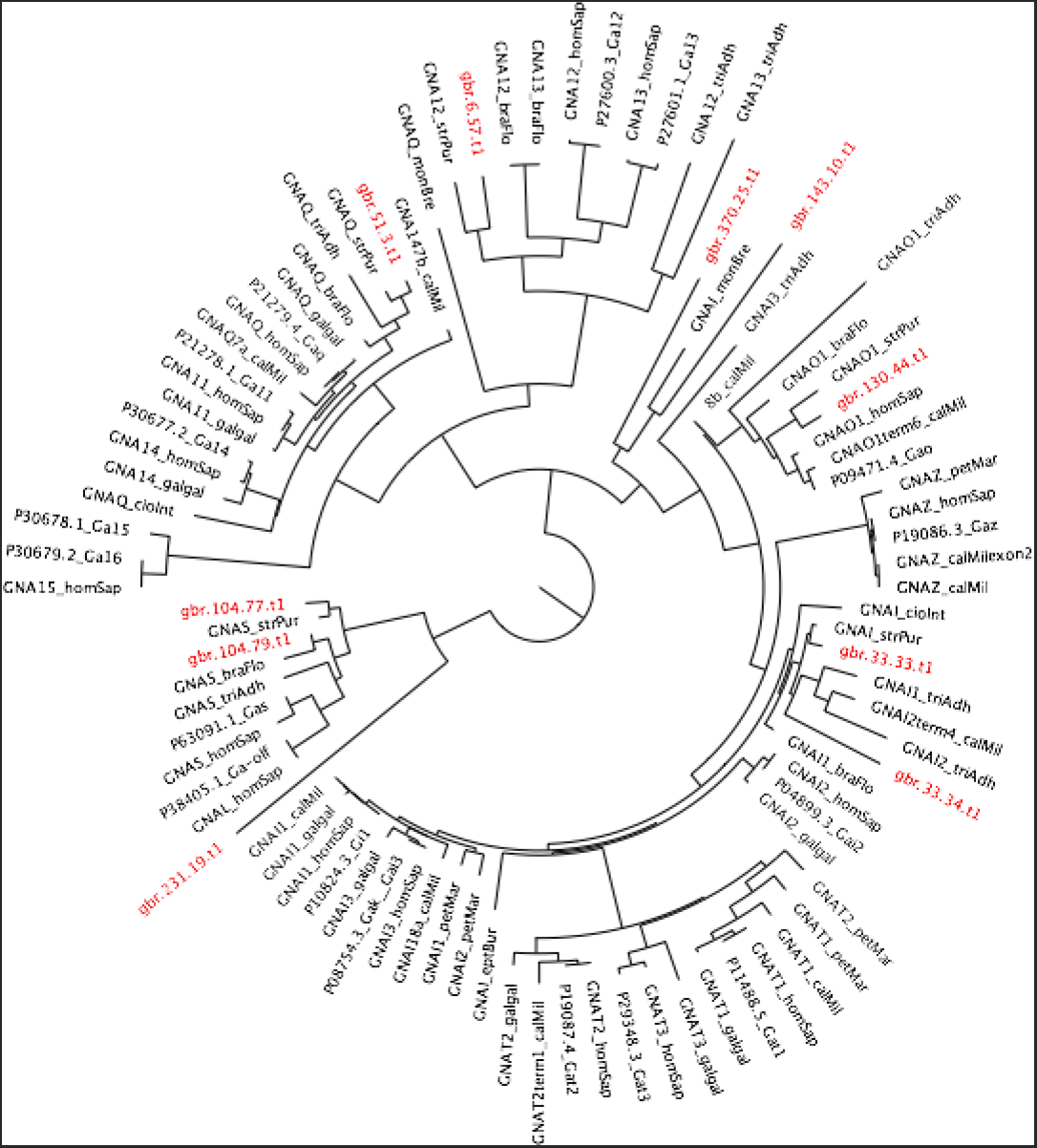
Phylogenomic tree of 92 G-protein alpha subunit sequences. *A. planci* sequences ID’s are highlighted in red. Of the 10 *A. planci* sequences 3 classified as Gα_s_, 1 as Gα_o_, 4 as Gα_i_, 1 as Gα_q_, and 1 as Gα_12_.

**Figure S3.**
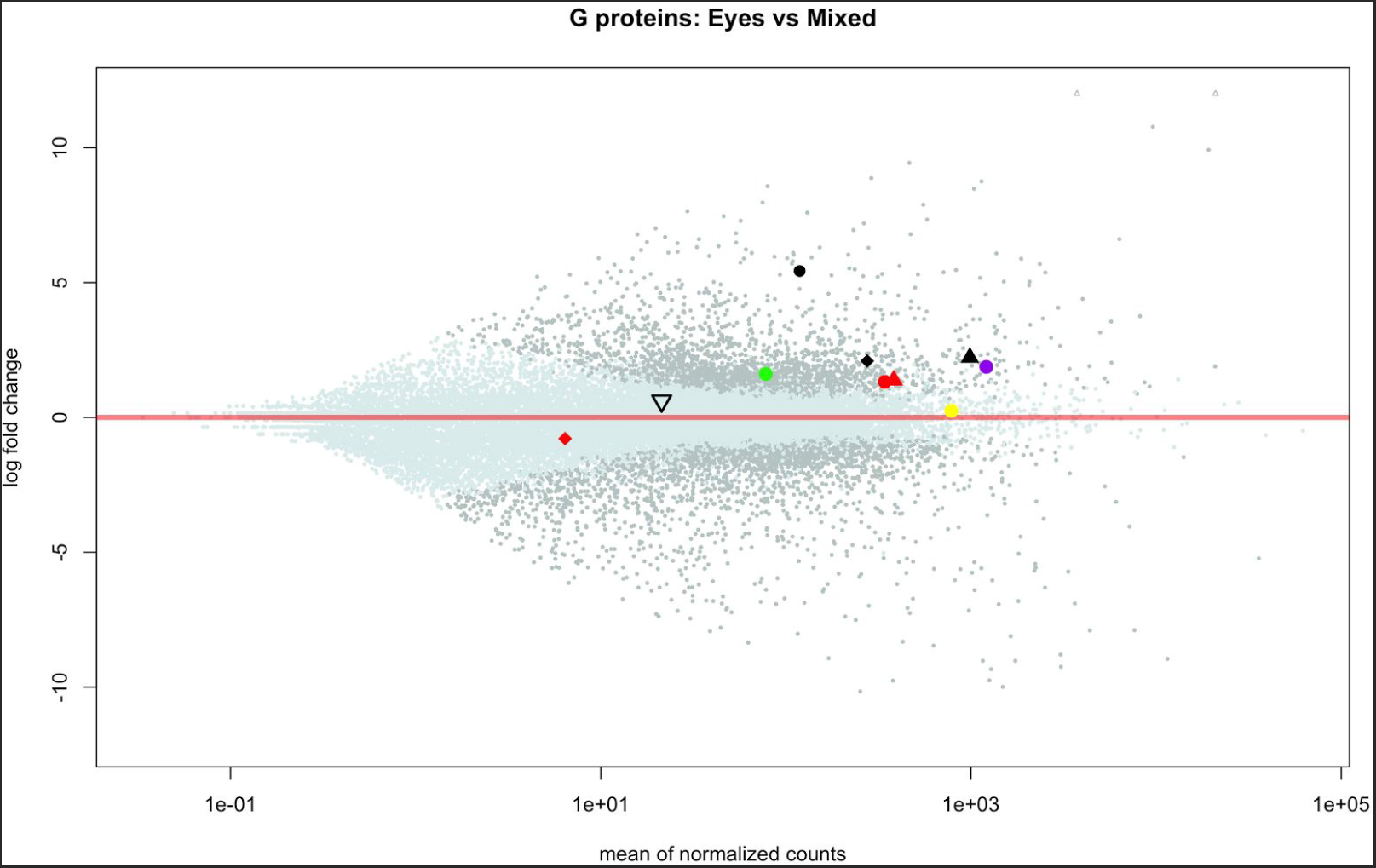
Differential gene expression in the *A. planci* eye samples compared to mixed tissues, with the G protein alpha subunits highlighted: 3 Gα_s_(red), 1 Gα_o_ (green), 4 Gα_i_ (black), 1 Gα_q_ (purple), and 1 Gα_12_ (yellow). All identified g protein alpha subunits with the exception of 1 Gα_s_ (gbr.231.19.t1), 1 Gα_i_(gbr.143.10.t1) and the Gα_12_are upregulated in the eyes of *A. planci* compared to the mixed tissue samples.

## Footnotes

1 COTS is thought to be divided into four separate species distinguished by bodies of water [23]-Red Sea, the Pacific, the Northern and the Southern Indian Ocean, reclassifying the Pacific ocean species as *Acanthaster solaris* [24]. However, for consistency with the released genome we will refer to the GBR species as *A. planci*.

